# MS2Lipid: a lipid subclass prediction program using machine learning and curated tandem mass spectral data

**DOI:** 10.1101/2024.05.16.594510

**Authors:** Nami Sakamoto, Takaki Oka, Yuki Matsuzawa, Kozo Nishida, Aya Hori, Makoto Arita, Hiroshi Tsugawa

## Abstract

Untargeted lipidomics using collision-induced dissociation-based tandem mass spectrometry (CID-MS/MS) is essential for biological and clinical applications. However, annotation confidence is still guaranteed by manual curation by analytical chemists, although various software tools have been developed for automatic spectral processing based on rule-based fragment annotations. In this study, we provide a novel machine learning model, MS2Lipid, for the prediction of lipid subclasses from MS/MS queries to provide an orthogonal decision of lipidomics software programs to determine the lipid subclass of ion features, in which a new descriptor, MCH (mode of carbon and hydrogen), was designed to increase the specificity of lipid subclasses in nominal mass resolution MS data. The model trained with 5,224 and 5,408 manually curated MS/MS spectra for the positive- and negative-ion modes mapped the query into one or several categories of 97 lipid subclasses, with an accuracy of 95.5% queries in the test set. Our program outperformed the CANOPUS ontology prediction program, providing correct annotations for 38.7% of the same test set. The program was further validated using various datasets from different machines and curators, and the average accuracy exceeded 87.4 %. Furthermore, the function of MS2Lipid was showcased by the annotation of novel esterified bile acids, whose abundance was significantly increased in obese patients in a human cohort study, suggesting that the machine learning model provides an independent criterion for lipid subclass classification, in addition to an environment for annotating lipid metabolites that have been previously unknown.

## Introduction

Lipids play vital roles in cellular membranes, signaling molecules, and energy storage in living organisms. Dysfunction in lipid metabolism is associated with human diseases^1^. The ClassyFire chemical classification program^2^ categorizes over 48,000 molecules as “lipids and lipid-like molecules,” as recorded in LIPID MAPS^3^ (as of January 6th, 2024). Mass spectrometry-based lipidomics techniques are currently used to elucidate the diversity and abundance profile of lipids. Untargeted lipidomics, utilizing liquid chromatography coupled with tandem mass spectrometry (LC-MS/MS) and electrospray ionization (ESI), efficiently provides the profile of 500 to 1000 lipid molecules from a single biospecimen. The structure of these lipids is elucidated by the pattern of product ion spectra generated by collision-induced dissociation (CID)^4^.

Numerous software programs support spectral mining for lipid structure elucidation^5^. Rule-based algorithms that confirm the presence of diagnostic ions to characterize lipid main classes, subclasses, and acyl chain compositions are commonly recommended for lipid structure description, guided by lipidomics standards initiative^4-6^. The term “lipid subclass” refers to a chemical class defined by the acyl chain linkage (e.g., ester, ether, and vinyl ether) in addition to the main class definition, such as phosphatidylcholine (PC) and phosphatidylethanolamine (PE). LIPID MAPS contains over 300 lipid subclass terms^3^. However, the current diagnostic tools are inadequate for determining lipid subclasses, as misannotations can occur due to contaminant ions from co-eluted lipids, in-source fragments, cluster ions, and spike noises generated by MS detectors^7^. Therefore, an orthogonal criterion is needed to improve annotation confidence in untargeted lipidomics.

Since the launch of the MS-DIAL 4 environment in 2020^8^, the program has been used in our laboratory for 82 projects involving 16,614 biological samples. Lipid annotations have been performed using a decision tree-based fragment annotation program for 117 lipid subclasses with a 1.5-minute retention time (RT) tolerance from reference RT values predicted by a machine learning method, as described in the previous report. The MS-DIAL program has provided lipid names for a total of 82,397 MS/MS spectral data in the alignment tables. The annotation results have been manually curated, with 39,871 spectral records labeled as “confidence” and 42,526 spectra labeled as a “mixture of several lipids” or “misannotation (false hit)”.

In this study, we developed a machine learning model called MS2Lipid using correctly labeled spectral records to predict lipid subclasses. Machine learning research on mass spectrometry data is actively conducted in metabolomics (including lipidomics) and proteomics fields^9,10^. In proteomics, there is a strong effort to construct machine learning models that use spectrum data obtained from biological samples to improve spectrum generation models and annotation accuracy^10^. However, in the field of metabolomics, most studies still primarily utilize standard spectra as training data. Our study is the first to employ manually curated lipid spectra derived from biological samples as training data in lipidomics. To evaluate the accuracy and precision of MS2Lipid, we used the CANOPUS program, which utilizes spectra records of authentic standard compounds, as a benchmark^11^. The MS2Lipid program was also evaluated on spectra containing at least two different lipids and on spectra where the lipid subclass origin was not trained in the machine learning process. Since the training dataset spectra were obtained from a single machine, we evaluated the scalability of MS2Lipid by using spectra from different machines and curators. Finally, we investigated the use of MS2Lipid for annotating previously unknown lipid metabolites in the reanalysis of a publicly available human cohort study^12^.

## Methods

### Mass spectral data for creating MS2lipid machine learning model

A total of 16,614 samples from 82 projects were analyzed using the same analytical method^13^ and processed in MS-DIAL 4, where 117 lipid subclasses could be characterized with the rule-based annotation program in combination with less than 1.5 min retention time tolerance matched with predicted retention times of lipids. All experimental data were acquired using reverse-phase liquid chromatography coupled with a SCIEX TripleTOF 6600 system, according to a previously described protocol^13^. Lipid structures were described using the MS/MS spectra obtained by data-dependent acquisition (MS/MS) in both positive- and negative-ion modes.

An experienced analytical chemist with more than 10 years of experience in analyzing lipidomic data curated the annotation results of the MS-DIAL. The annotated spectra in the alignment files were manually evaluated. In total, 82,397 annotated spectra from 164 alignment result files derived from 82 positive- and 82 negative-ion mode data were curated. The annotation is labeled as “incorrect (false hit)” in the cases that the peak shape looks noisy (signal-to-noise ratio approximately less than 10) or the diagnostic ions are matched with barcode (noise) ions. This data curation process was not conducted using a systematic method; instead, the final judgement for “correct,” “mix,” and “false” labelings was made based on the knowledge and experience of the skilled technician. It is important to note that the labeling work was not performed for all annotated peaks. In our study, lipid quantification was performed using a representative adduct form that is independent of each lipid subclass independently^8^. For example, phosphatidylcholine (PC) is commonly detected in adduct ion forms such as [M+H]^+^, [M+Na]^+^, and [M+CH_3_COO]^-^, whereas only [M+CH_3_COO]^-^ is used to quantify lipid metabolites. Therefore, the annotated MS/MS spectra derived from PC’s [M+H]^+^ and [M+Na]^+^ PCs are often not labeled in routine work. As a result, machine learning of the spectra for adduct ion forms not targeted by our group may be insufficient.

Total 39,871 spectra encompassing 4,100 metabolites of 116 lipid subclasses were labeled as “correct” (**Supplementary Data1**, on marked as “ok” in comment column). Spectral records of lipid molecules with fewer than four record entries in either ion mode (positive- or negative-ion mode) were excluded. The spectral records of free fatty acids (FA) and *N*-acyl ethanolamine (NAE) were excluded because the annotations were performed only based on the information of retention time and *m/z* values due to the lack of diagnostic criteria in the MS/MS spectra. Moreover, lipid spectral records having “others” as the ontology term were excluded: in the MS-DIAL lipidomics project, standard compound spectral library-based peak annotation based on the records of MassBank, GNPS, and NIST is also executed when the spectrum is not characterized by the rule-based lipid annotation pipeline. Finally, 16,614 spectral records of 3,944 unique lipid molecules across 97 lipid classes were used for the machine learning (**Supplementary Data2**). It contained 8,451 ESI(+)-MS/MS spectra of 2,559 unique lipids and 8,163 ESI(-)-MS/MS spectra of 2,048 unique lipids.

### Spectral data for the validation of MS2Lipid

Annotated tandem mass spectral data from 31 projects were downloaded from the RIKEN LIPIDOMICS website (http://prime.psc.riken.jp/menta.cgi/lipidomics/index) (**Supplementary Data3**). These spectral data were curated by different analysts from the training data used in this study. Lipidomic data were obtained using various MS machines, including the SCIEX TripleTOF 5600+ (index 18 to 28), Waters XevoG2 QTOF (index 78), ThermoFisher Q-Exactive Plus (index 79), Agilent 6546 QTOF system (index 80), SCIEX TripleTOF 6600 using SWATH-DIA (index 81), and Bruker timsTOF Pro (index 83 and 84). In addition, several studies (indexes 1 to 11) were obtained using the same analytical instrument (SCIEX TripleTOF 6600) but curated by different analysts.

### Data preprocessing for machine learning

Product ions (PLs) ranging from *m/z* 70–1250 were used, and the *m/z* values were rounded to one decimal place. The centroid peak heights in the bins were summed. In addition to the vector of the product ions, neutral loss (NL) information from the precursor *m/z* was used for the variables. The NL value was defined as the mass difference between the precursor *m/z* and product ion *m/z* values. The NL values ranging from 0 to 10 were excluded. The intensity of the NL vector was prepared in the same manner as that of the PL vector. The *m/z* and NL values with zero standard deviations in the dataset were excluded. Peak intensities were normalized, and the base peak intensity was standardized to 1. In addition to the PL and NL vectors, two bins were prepared. The first feature was a binary value representing odd- or even-round precursor *m/z* values. The second feature value was designed to be the same within the same lipid subclass and differentiate lipid subclasses with similar spectral patterns. For example, the *m/z* values of the precursor ions PC 33:1 (C_41_H_80_NO_8_P) and PC O-34:1 (C_42_H_84_NO_7_P) were 804.576 and 804.612, respectively, where the rounded precursor values were the same and the MS/MS spectral patterns were indistinguishable. Because the exact mass difference between PC 33:1 and PC O-34:1 arises from the exact masses of CH_4_ and O, a new property value, termed the mode of carbon and hydrogen (MCH) value, was defined by the following equation (**Supplementary Figure 1 and Supplementary Table1**).

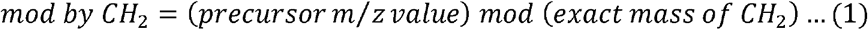

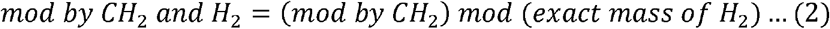

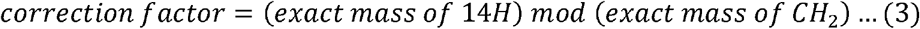

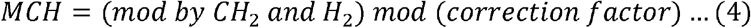

The MCH value was designed to provide a consistent value among molecules of each lipid subclass (see the evidence in **Supplementary Note 1**), and was calculated based on the modulus operation to calculate the remainder in the division process. The remainder of the precursor *m/z* value based on the exact mass of CH_2_ was calculated (Equation 1). Furthermore, the reminder value was further dedicated to the modulus operation by the exact mass of H_2_ where the differences in acyl chain length and double bond number were canceled with the condition of fewer than six double bonds (equation 2). To provide a consistent value that was not affected by the double bond number, the correction factor was calculated (Equation 3). A correction factor is applied to calculate the remainder generated using Equation 2 (Equation 4).

### Selection of machine learning model to predict lipid subclass

Machine learning was performed using Python version 3.11.6. One-fifth (20%) of the dataset was used as the test set. Of the remaining data (80%), four-fifths (64%) were used for the training set, and one-fifth (16%) was used for the validation set. In this study, support vector machine (SVM), k-nearest neighbor (KNN), random forest (RF), and deep neural network were investigated. Parameter optimizations for SVM, KNN, and RF were performed using scikit-learn (https://scikit-learn.org/stable/#), and the optimal parameters in the deep neural network (dNN) were determined using the search function of the Keras Tuner Library (https://keras.io/keras_tuner/) using the training and validation datasets. The performance was evaluated using accuracy, recall, precision, and F1 scores. For each method, the following parameters were optimized by a random search method, where accuracy was used for parameter selection: SVM, regularization parameters, kernel, decision function; KNN, number of neighbors, weights, distance function; RF, number of neighbors, criterion function, maximum tree depth; and dNN, number of layers, thickness of each layer, batch correction method, learning rate, and dropout rate. The source code for the model selection is available on the GitHub website (https://github.com/systemsomicslab/ms2lipid/tree/main/notebooks). Although the performances of the optimized SVM, KNN, and dNN models were mostly equal, the dNN model was selected in this study as the best model because of its high accuracy.

### MS2Lipid model using deep neutral network with curated spectral records

A deep learning model of MS2Lipid was constructed for the ESI(+)-MS/MS and ESI(+)-MS/MS spectral records. In the positive ion mode, the number of input and output features were 2,077 variables and 63 lipid subclasses, respectively. A rectified linear unit (ReLU) was used as the activation function. The softmax activation function was used in the output layer. The number of hidden layers was restricted to 2–10 layers. The dropout rate was also restricted from 0.1 to 0.5. Consequently, the tuner provides 1,664, 2,304, and 3072 nodes for the first, second, and third layers, respectively, with a 50% dropout probability. In the negative-ion mode, the numbers of input and output features were 2,816 variables and 69 lipid subclasses, respectively. The tuner offered 3,958 and 2,176 nodes for the first and second hidden layers, respectively, with a 40% dropout probability. The output layer represents the probability of the lipid ontology classification. The lipid subclass with the highest probability value was used as a representative candidate for the predicted lipid subclasses. In this study, if the representative lipid subclass is matched with the correct lipid subclass, the result becomes “correct.” If the correct lipid subclass is listed in the predicted lipid candidates where the probability value exceeds 1%, the result is defined as “listed in candidates.”

### Using Shapley Additive exPlanations (SHAP) for important variables interpretation

The important variables for deep neural network machine learners were evaluated using the SHAP values^14^. Because of computational cost limitations, 100 spectral records containing 10 unique lipid subclasses, each containing 10 molecules, were incorporated into the SHAP function using the Python package (https://github.com/shap/shap). Spectral records with the top 10 precursor ion intensities were selected for each lipid subclass.

### Evaluation of CANOPUS for the benchmark of MS2Lipid

The spectral records of the test datasets were analyzed using the CANOPUS ontology prediction program implemented in SIRIUS^15^. Ontology terms followed the definition of ClassyFire^2^. The records were converted to the “. ms” format files as defined in the program. SIRIUS version 5.8.2 was utilized. To compare the results of MS2Lipid and CANOPUS, the subclass definition of LipidMAPS/MS-DIAL was converted to the definition of ClassyFire “class” definition (**Supplementary Table 2**). In this study, 13 ClassyFire classes were identified: fatty acids and conjugates, glycerophosphocholines, glycerophosphoethanolamines, glycerophosphoinositols, glycerophosphates, glycerophosphoserines, glycerophosphoglycerol, phosphosphingolipids, glycosphingolipids, ceramides, diacylglycerols, monoacylglycerols, and triacylglycerol.

### Reanalysis of a publicly available human feces lipidomics data

A publicly available LC-ESI(+)-MS/MS- and LC-ESI(-)-MS/MS data sets from a human cohort study where the associations between insulin resistances and microbiome metabolisms in 306 individuals were downloaded from RIKEN DROPMet website (index DM0037, http://prime.psc.riken.jp/menta.cgi/prime/drop_index). The LC-MS data were analyzed using MS-DIAL 5^16^ with the parameter set available in **Supplementary Table 3**, where the retention time correction function was used to improve the peak alignment process (**Supplementary Table 4**). The alignment results for the positive and negative ion modes are presented in **Supplementary Data 4**. For the spectrum network analysis, the MS/MS spectral similarity was calculated using the Bonanza scoring system of MS-DIAL^17^ where the parameters were set as follows: minimum *m/z*, 70; relative abundance cutoff, 1; absolute abundance cutoff, 12; mass tolerance, 0.025; minimum peak match, 2; match threshold, 0.7 (70%); and maximum edge number per node = 20. Peak features where the average signal-to-noise ratio among samples was more than 10 were visualized using Cytoscape version 3.10.1 (https://cytoscape.org/). For the correlation analysis between the lipidome and microbiome data, 26 bacterial genera with less than 50% of samples having zero counts were used. The relationship between lipids and the gut microbiota was evaluated using Spearman’s correlation.

## Results

### Vector construction for machine learning using lipid tandem mass spectra

We utilized 82 studies comprising a total of 82,397 spectra for vector construction. The MS-DIAL program^9^ was employed for peak picking, annotation, and peak alignment. After the curation for annotated peaks (see **Methods**), our study employed a total of 16,614 spectra from 3,944 unique lipids belonging to 97 subclasses. This included 8,451 ESI(+)-MS/MS spectra of 2,559 unique lipids and 8,163 ESI(-)-MS/MS spectra of 2,048 unique lipids. The mass spectrum is represented as a vector through the following procedure (**Figure 1**) (also see **Methods**). The high-resolution mass values were converted to nominal mass values using a simple rounding method. In addition to the vector of product ions, the neutral loss (NL) from the precursor *m/z* value was also included as a variable. Furthermore, two additional descriptors were created for this study. One descriptor indicates whether the rounded precursor *m/z* value is even or odd. This binary value is particularly important for distinguishing between phosphatidylcholine (PC) and sphingomyelin (SM), as their product ion spectrum patterns are similar in positive ion mode due to the highly sensitive product ion of *m/z* 184 related to the phosphocholine polar head group. The second descriptor is designed to distinguish between two independent subclasses that have very similar spectra but can be distinguished by their high-resolution *m/z* values **(Supplementary Figure 1 and Supplementary Table 1)**. An exemplary case is the differentiation between PC and ether-linked PC. For instance, PC 15:0_18:1 and PC O-16:0_18:1 have molecular formulas and exact mass values of C_41_H_80_NO_8_P and 745.562, and C_42_H_84_NO_7_P and 745.599, respectively, resulting in a 37 mDa mass difference. These metabolites can be distinguished using high-resolution MS. Therefore, we created a value called “mod by carbon-hydrogen mass values (MCH-value),” which provides a consistent value for each lipid subclass **(Supplementary Note 1)**.

**Figure 1.**
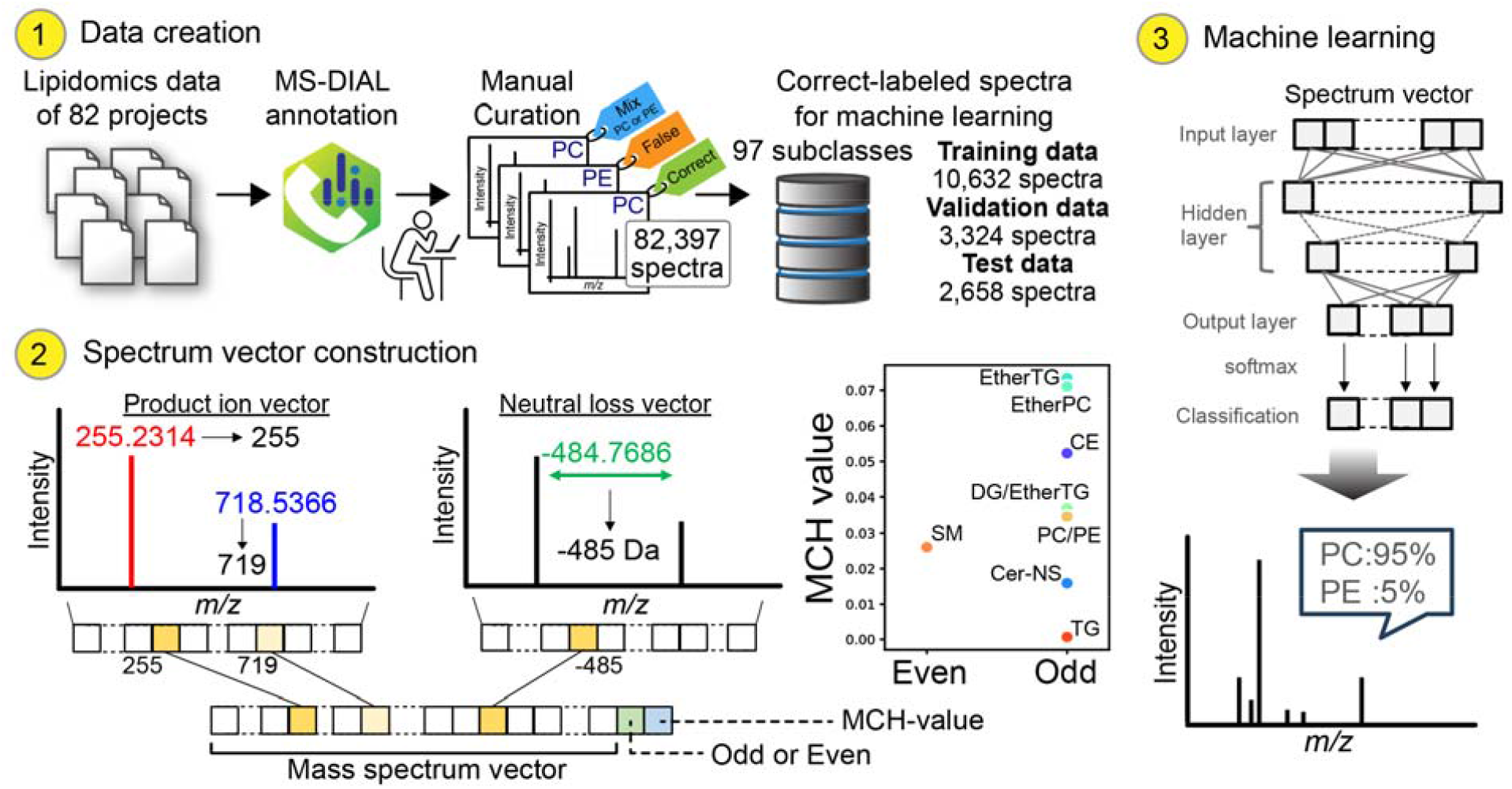
Workflow to create MS2Lipid machine learning model. In data creation step, the spectral data were derived from 82 projects. The lipidomics data were analyzed by MS-DIAL, providing lipid annotation for 82,397 spectra that include positive- and negative ion modes. The original annotation was manually curated, and labeled as correct, mix, or false hit. The correct-labeled data of 16,614 spectra were divided into training, validation, and test sets. In spectrum vector construction step, the MS/MS spectrum is represented by the array of product ions and neutral losses whose high-resolution mass value is converted into nominal mass. In addition, two descriptors, “odd or even” of precursor nominal mass value and “MCH-value” calculated from the accurate precursor *m/z* value. The descriptors for cholesteryl ester (CE), ceramide containing sphingosine and normal fatty acid (Cer-NS), diacylglycerol (DG), ether-linked DG (Ether DG), phosphatidylcholine (PC), ether-linked PC (Ether PC), triacylglycerol (TG), ether-linked TG (ether TG), phosphatidylethanolamine (PE), and sphingomyelin (SM) were described. The vectors were used as the input for machine learning models. The output shows the probability ratio of lipid subclass classifications.

### Selection and optimization of machine learning models

We evaluated four machine learning methods: support vector machine (SVM), k-nearest neighbor (KNN), random forest, and deep neural networks (**Figure 2a**) (see **Methods** for the parameter optimizations). The results indicated that the model using a deep neural network offered the best accuracy in both ion modes. We evaluated the optimized model using the test dataset, in which the lipid subclass with the highest probability value was defined as the representative candidate. In the case that the correct lipid subclass was predicted by the probability of >1%, the result was defined as “listed in candidates” (**Figure 2b**). The result showed that the correct lipid subclass was predicted with accuracies of 96.1% and 94.9% for the positive and negative ion mode spectra, respectively. Moreover, the correct lipid subclass was listed as a candidate for more than 98% of the queries.

**Figure 2.**
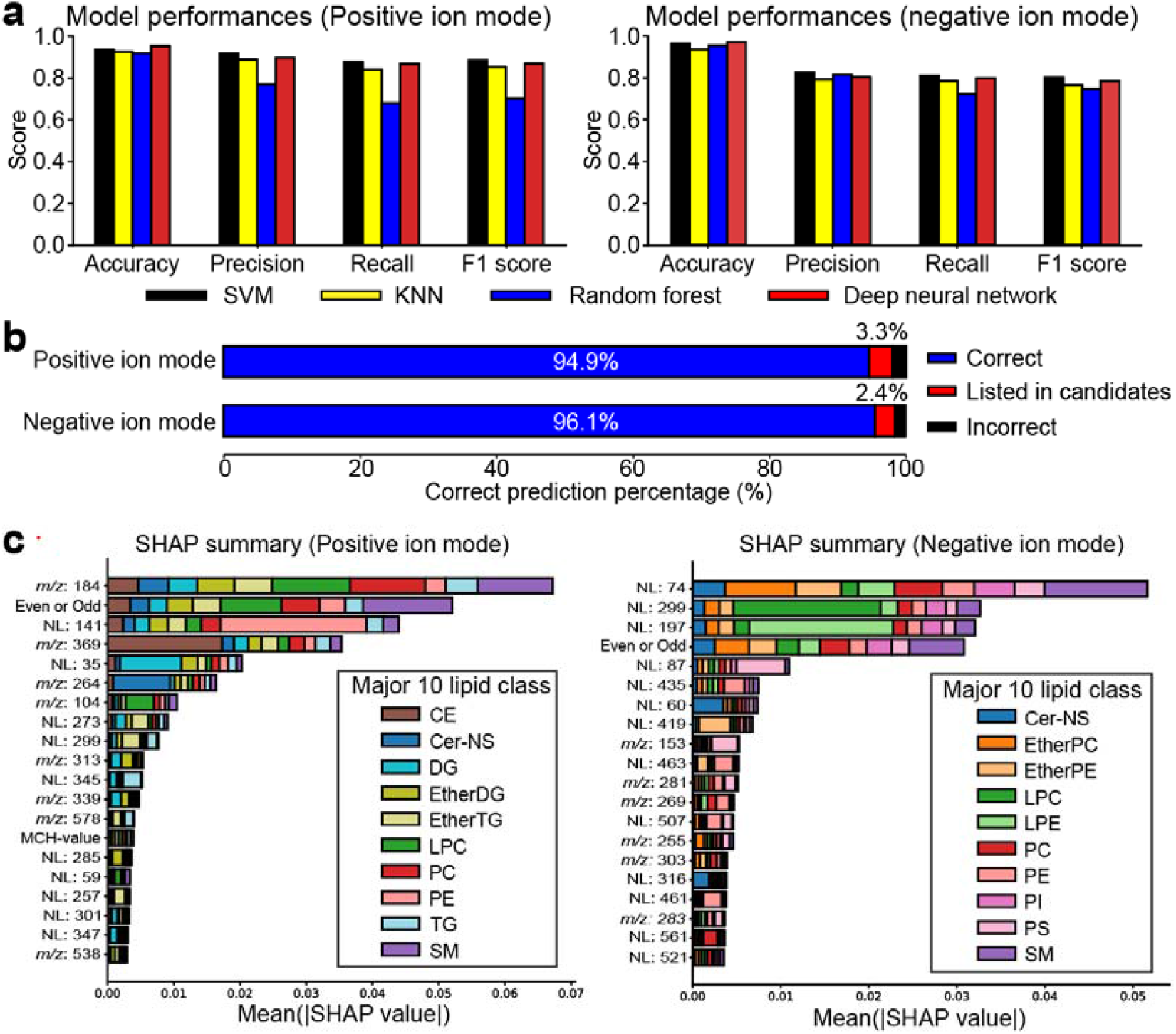
Evaluation of machine learning models. (a) Accuracy, precision, recall, and F1-score in each model. Support vector machine (SVM), k-nearest neighbor (KNN), random forest, and deep neural network were evaluated. (b) Prediction accuracy in the deep neural network model. The upper- and lower panels correspond to the results in positive- and negative ion modes, respectively. The output becomes correct if the predicted lipid subclass is equal to the correct label, and the result becomes “listed in candidates” if the correct label exists in the candidate list. (c) The shapley additive explanation (SHAP) scores to investigate important descriptors. The top 20 SHAP features for predicting 10 major lipid subclasses were described. NL, neutral loss; phosphatidylethanolamine; PI, phosphatidylinositol; PS, phosphatidylserine; EtherPE, ether-linked PE; LPC, lysoPC; LPE, lysoPE.

The shapley additive explanation (SHAP) score was calculated to investigate the important descriptors for lipid subclass classification (**Figure 2c**)^14^. Owing to the limitations of the SHAP calculation cost, the important features for predicting the 10 lipid subclasses were evaluated for each ion mode. The neutral losses (NLs) of 74, 299, 197, and 87 Da were extracted as important features in the negative ion mode. The NLs denote for the loss of acetic acid plus methyl (C_3_H_6_O_2_, NL of 74.037 Da) in PC and SM, the loss of acetic acid plus choline alfoscerate (C_10_H_22_NO_7_P, NL of 299.113 Da) in lyso PC (LPC), the loss of ethanolamine glycerophosphate (C_5_H_14_NO_5_P, NL of 199.061 Da) observed in lyso PE (LPE), and the loss of serine polar head (C_3_H_5_NO_2_, NL of 87.032 Da) observed in phosphatidylserine (PS). The product ions of *m/z* 184, NL of 141 Da, and *m/z* 369 were extracted as important features in the positive ion mode, which denote the phosphocholine polar head (C_5_H_15_NO_4_P^+^, *m/z* 184.073), phosphoethanolamine polar head (C_2_H_8_NO_4_P, NL of 141.019 Da), and cholesterol aglycone (C_27_H_45_^+^, *m/z* 369.349), respectively. Moreover, the result indicated that the descriptor of “odd or even” was expected as the important feature in both positive- and negative ion modes, which contributed to the judgement of sphingolipids (SM) and PC as expected. In addition, the descriptor of “MCH-value” was extracted as the important feature (totesp 14) in positive ion mode. These SHAP results indicate that the MS2Lipid model recognizes diagnostic ions that have been utilized in the lipidomics community to predict lipid subclasses. Here, the accuracy of MS2Lipid for the test set was 95.5% (**Supplementary Table 5**).

### The performance comparisons between MS2Lipid and CANOPUS

We used CANOPUS^12^ as the benchmark for the MS2Lipid program, which classifies product ion spectra into chemical classes defined by ClassyFire. The same test set was used for both programs. It’s important to note that CANOPUS predicts all chemical classes, including lipids, from the query spectrum. The model was trained using authentic standard-derived spectral libraries such as GNPS and MassBank. However, the spectral records of lipids in these libraries are smaller compared to our training set. Therefore, the following comparison reflects the lipid diversity in the training sets rather than the model building process. For example, our training data set includes 5,224 spectra from 1,408 molecules of glycerophosphocholines, glycerophosphoethanolamines, glycerophosphoinositols, glycerophosphates, glycerophosphoserines, and glycerophosphoglycerols. In contrast, GNPS (GNPS-LIBRARY as of January 25, 2024) and MassBank (version 2023.11) databases have 60 spectra from 43 molecules and 2,579 spectra from 558 molecules, respectively^19,20^. Additionally, our training set includes 1,301 spectra from 456 triacylglycerols (TGs), while GNPS and MassBank databases have 7 spectra from 5 molecules and 65 spectra from 17 molecules, respectively (**Supplementary Table 6**).

Since our model generates lipid ontologies defined by LIPID MAPS or MS-DIAL, the output of the MS2Lipid program was converted to the compound class defined by ClassyFire. The accuracy of the MS2Lipid program in predicting the ClassyFire “class” level exceeded 99.0%. CANOPUS did not generate output for half of the spectral queries because it provides predicted results only for compounds with a probability above 50% and primarily supports computation for compounds less than 600 Da (https://boecker-lab.github.io/docs.sirius.github.io/). CANOPUS supported only 38.7% of the spectral queries. However, the MS2Lipid program outperformed CANOPUS in prediction accuracy (**Supplementary Table 7**), with CANOPUS achieving 91.8% accuracy for the queries that CANOPUS could handle (**Figure 3**). This result indicates that in lipid spectrum machine learning research, it is necessary to accumulate both standard spectra and spectral information from biological samples to build a highly accurate learning model.

**Figure 3.**
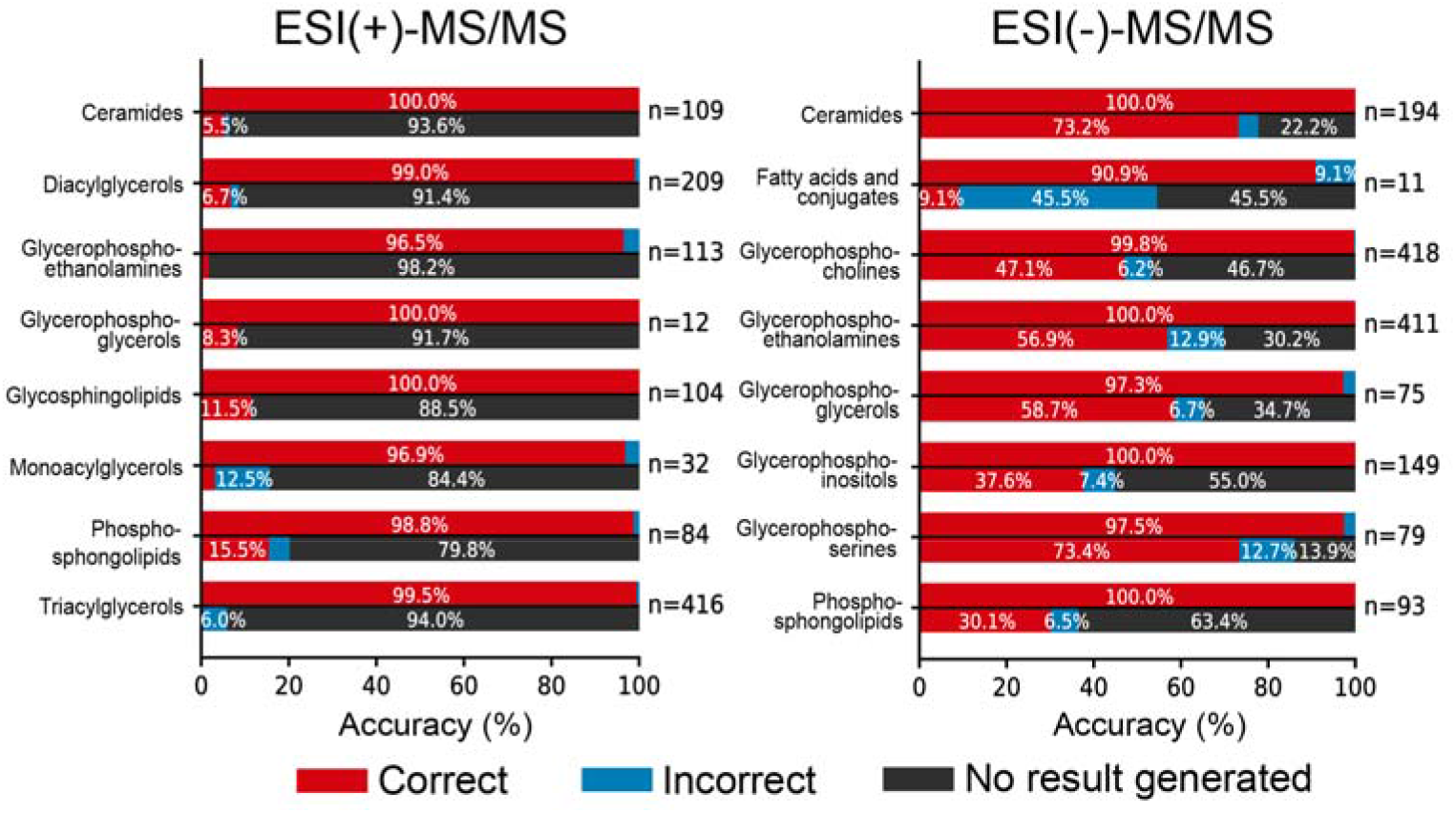
Comparison of MS2Lipid with CANOPUS. The left and right panels show the results of positive- and negative ion mode, respectively. In each lipid class, the upper- and lower panels are the results of MS2Lipid and CANOPUS, respectively. The accuracy (%) of ClassyFire class level ontology prediction was described. The number of molecules in each ontology was also shown.

### Evaluation of robustness and scalability of MS2Lipid by using various spectral data

We evaluated the robustness of MS2Lipid for spectra obtained using various machines and curated by various analysts. The data resources were downloaded from the RIKEN LIPIDOMICS website (http://prime.psc.riken.jp/menta.cgi/lipidomics/index), in which the spectral data from Waters, Bruker, Thermo Fisher, Agilent, and SCIEX were available. The spectral data in our training dataset were not included in the data resources. First, the MS/MS spectral data acquired by the same machine and curated by different analysts were imported into the MS2Lipid program, resulting in 94.5% and 81.4% accuracy for the positive and negative ion modes, respectively (**Figure 4a**). This is an important result because different data curators may offer different results, which reminds us of developing a machine-learning model that provides an orthogonal decision objectively, regardless of the data analysts. Next, we applied the MS2Lipid program to the spectra acquired using different curators and MS techniques (**Figure 4b**). While the accuracies ranged from 71% to 98%, the accuracy for Waters XevoG2 QTOF in the negative ion mode was 19.4%. This was due to the difference in solvent conditions, where ammonium formate-generating formate adduct form [M+HCOO]^-^ for neutral or cationic molecules was used in the experiment using Waters XevoG2 QTOF, while ammonium acetate-generating acetate adduct form [M+CH_3_COO]^-^ was used for the others, including our training data set, which results in a 14 Da mass shift in the product ion patterns. Because the neutral loss feature of 74 Da is the diagnostic criterion for PC and SM classification, all test queries for PC and SM lipids were misclassified using the MS2Lipid program. When lipid subclasses ionized with formate or acetate were excluded from the test dataset in the Waters XevoG2 QTOF spectra, the accuracy was 100%. Consequently, our results suggest that accurate predictions require metadata describing the details of the analytical conditions used. Finally, we demonstrated how the MS2Lipid program handled the product ion spectra derived from the co-eluted lipids (**Figure 4c**). An MS/MS spectrum considered to be a mixture of PC and PE molecules because of the presence of *m/z* 184 for the PC motif and an NL of 141 for the PE motif is shown. The MS2Lipid program provided the probability as the results in which the mixed spectra were predicted as “54.8% PE and 19.4% PC”, while the MS-DIAL program only offers either PC or PE as a representative candidate based on the similarity score: the percentage does not mean the abundance ratio of two metabolites.

**Figure 4.**
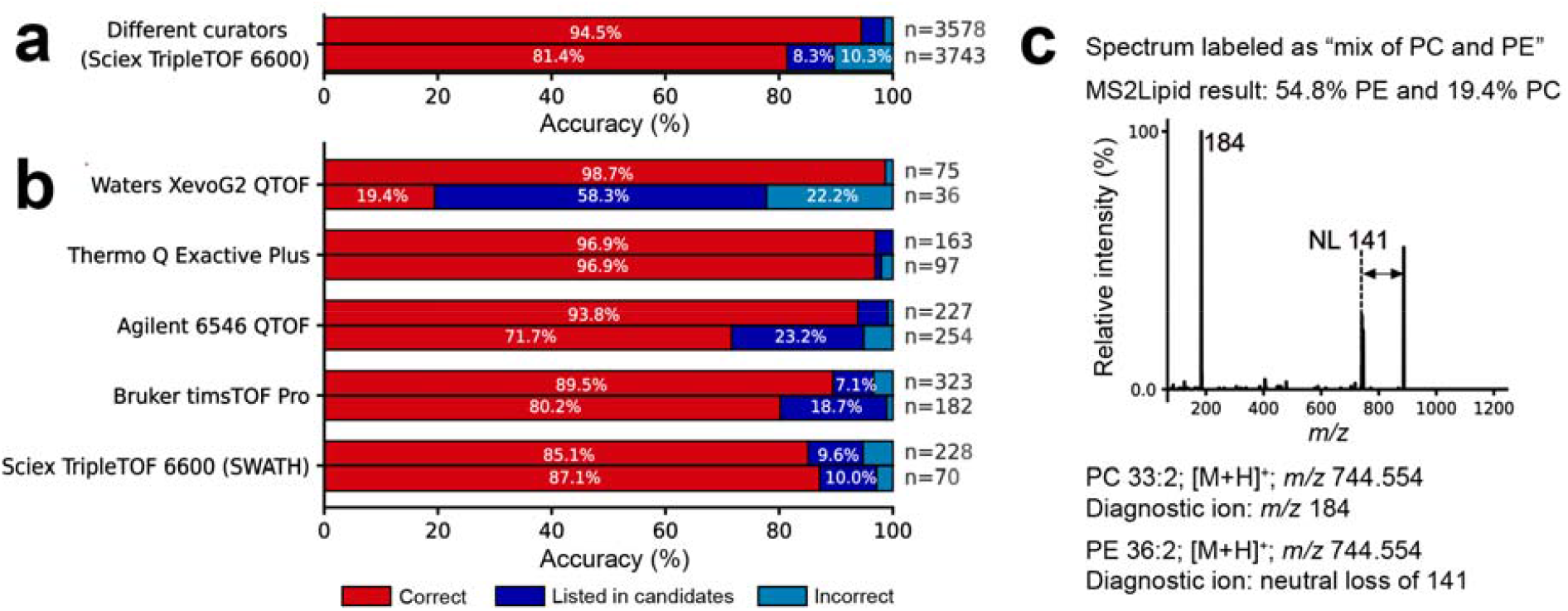
Evaluation of robustness and scalability of MS2Lipid. (a) Validation using the spectral data by different curators with the completely same MS machine. (b) Validation using the spectral records obtained by different instruments and different curators. The outputs are categorized into three cases: if the generated label of MS2Lipid is the same as the query’s label, the result becomes “correct”; while the labels are not matched, the result becomes “listed in candidates” if the correct name is listed in the candidates predicted with more than 1% probability; and for the others, the result becomes “incorrect”. (**c**) MS2Lipid result for predicting the mixed spectra of phosphatidylcholine (PC) and phosphatidylethanolamine (PE).

### Application of MS2Lipid for annotating lipids that have previously been unknowns in a human cohort analyzing the fecal lipidome

The MS2Lipid program was used to annotate the novel lipid molecules. We reanalyzed a publicly available LC-ESI(+)-MS/MS- and LC-ESI(-)-MS/MS data sets from a human cohort study where the associations between insulin resistances and microbiome metabolisms in 306 individuals have been investigated^12^. The MS-DIAL program generated 22,980 and 12,742 unique peak features in ESI(+)- and ESI(-) data sets, respectively, in which the MS/MS spectra for 18,497- and 10,224 features (precursor ions) were assigned. Of these, 4,135 of ESI (+) and 2,967 of ESI(-) spectra were characterized by the MS-DIAL rule-based annotation algorithms in combination with the predicted retention time information.

First, we applied the MS2Lipid program to the annotated MS/MS spectra using MS-DIAL. As a result, 54.1% and 48.4% of peaks (if considered listed as candidates; 67.6% and 60.6%) for ESI(+)- and ESI(-)-MS/MS spectra, respectively, were predicted with the same lipid ontology, where the lipid subclasses supported by the current MS2Lipid program were evaluated. This indicated that half of the MS/MS spectra characterized by MS-DIAL were not assigned by the MS2Lipid program. In contrast to MS-DIAL, which estimates lipid structures based on the presence or absence of diagnostic ions without considering the effects of noise or contaminated ions, MS2Lipid determines lipid classes by considering the presence of noise and contaminated ions, reflecting the discretion of experienced analysts (**Figure 2a**). This suggests that MS2Lipid performs more conservative annotations, which could explain why more than half of the spectra analyzed by MS-DIAL were not supported by MS2lipid.

Next, we applied the MS2Lipid program to the unknown spectra, in which 14,272 peaks with MS/MS spectral information were labeled as unknown in MS-DIAL. The molecular spectrum network was constructed (**Figure 5a**). The top-hit annotations by MS-DIAL and MS2Lipid were mapped onto the network, and the results were reflected by node fill and border colors, respectively. Distinct clusters based on major lipid subclasses were created in the molecular spectrum network, and we found a cluster where 78% of the nodes were not characterized by the MS-DIAL program. The cluster contains a subclass of bile acid esters assigned as steryl ester (SE) 24:1;O4/FA, whose well-known structural backbone is deoxycholic acid (DCA) and its isoform (isoDCA), which is the second bile acid biosynthesized by bacteria^18^. Hereafter, we define the bile acid ester assigned by SE 24:1;O4/FA as “DCAE” implying deoxycholic acid ester. Interestingly, most of the unknown peak features were predicted to be DCAE, other sterylesters, DCA, and *N*-acyl glycine. The existence of the term *N*-acyl glycine in addition to DCAE enabled us to elucidate unknown molecules such as glycine-conjugated deoxycholic acid ester (GDCAE), which contains *m/z* 414 as a unique ion (**Figure 5a**). Because the stereochemistry of the sterol backbone cannot be characterized by the CID-MS/MS technique, the characterization of the molecules with *m/z* 414 was assigned as SE 24:1;O4;G/FA in our study, where the nomenclature follows the definition of LIPID MAPS. It has been reported that the bile acid of isoDCA is efficiently converted to its ester form^19^. Furthermore, neither SE 24:1;O4;G/FA nor SE 24:1;O4/FA were detected in our previous study analyzing mouse feces^8,20^, and isoDCA was less abundant in mouse feces than in human feces^18^. Furthermore, the ester form of ST 24:1;O5, implying cholic acid, was not detected in human feces, which was also true in mouse feces. This indirect evidence enabled us to expect the structure to be a glycine-conjugated isodeoxycholic acid ester, although its complete structure should be confirmed using an authentic standard in the future.

**Figure 5.**
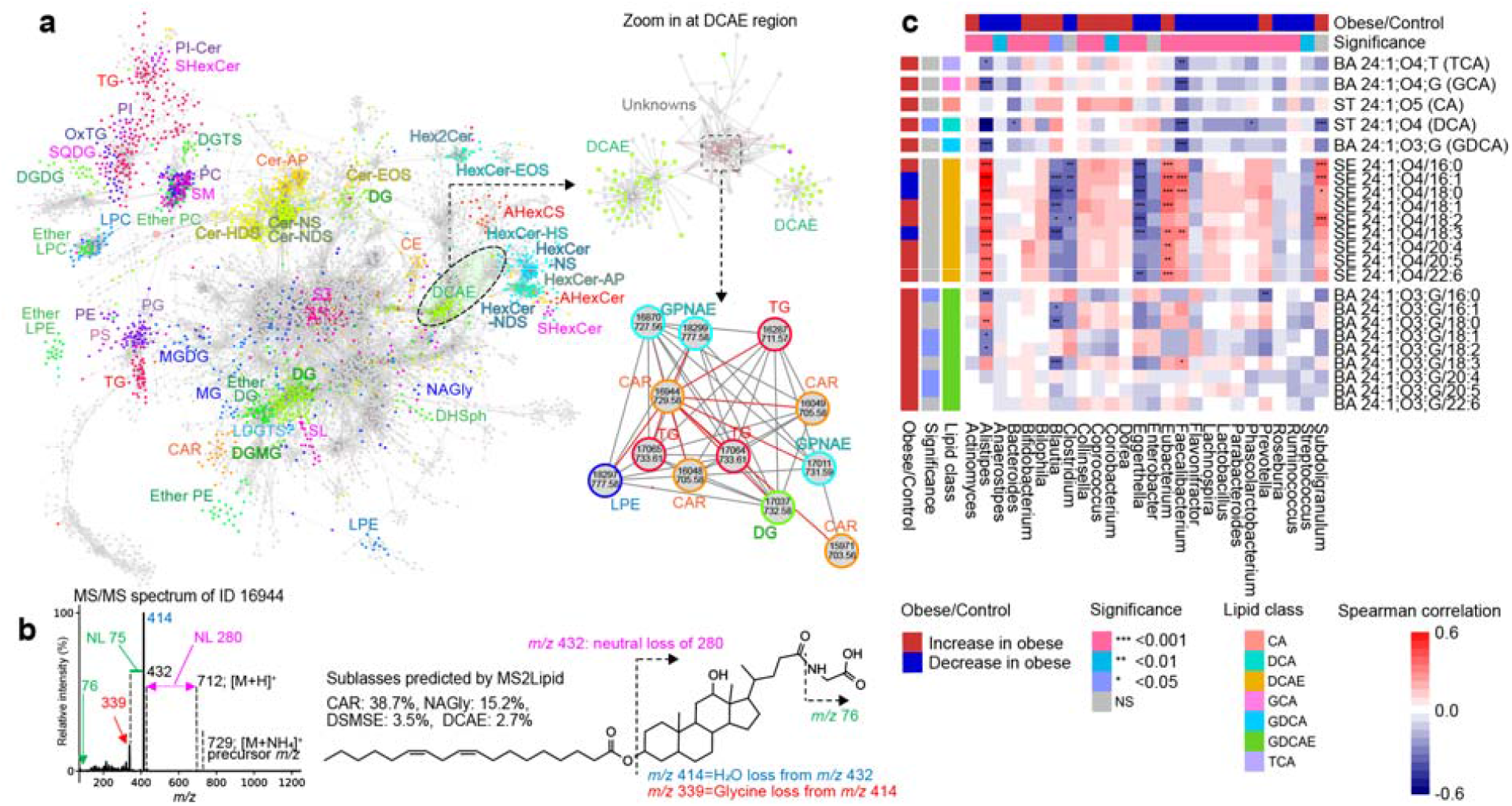
Mapping MS-DIAL and MS2Lipid ontology terms into a molecular spectrum network. (**a**) Molecular spectrum networking from peak features of positive ion mode in a human cohort study analyzing fecal samples. The node fill- and border colors represent the annotation of MS-DIAL and MS2Lipid, respectively. (b)The tandem mass spectrum of newly characterized bile acid ester and its plausible structure were described. The features of *m/z* 75 and neutral loss (NL) of 76 originate from the glycine structure. The features of *m/z* 339, NL 260, and *m/z* 414 are derived from the lipid subclasses of deoxycholic acid, FA 18:2-containing lipid, and the glycolic acid structure, respectively. (**c**) Heatmap analysis between bacteria and lipid molecules. Statistical testing for each lipid or bacteria in between control and obese individuals was performed. Significance levels in spearman correlation denoted as *: 0.05, **: 0.01, ***: 0.001. CA; Cholic acid, DCA; Deoxycholic acid, DCAE; Esterified deoxycholic acid, GCA; Glycocholic acid, GDCA; Glycodeoxycholic acid, GDCAE; Esterified glycodeoxycholic acid, TCA; Taurocholic acid.

We explored the relationship between the amount of GDCAE molecules and the clinical diagnosis of obese and healthy individuals (**Figure 5b**). The abundance of glycine-conjugated molecules, including FA 16:0, 18:1, 18:2, 20:4, and 20:5, was significantly increased in obese individuals. Furthermore, correlation analysis between bile acids and bacteria revealed that the abundance of DCAE was highly correlated with *Alistipes, Eubacterium, Faecalibacterium*, and *Subdoligranulum*, whereas the abundance of GDCAE molecules was negatively correlated with *Alistipes, Blautia*, and *Prevotella*. Given that the highly abundant molecules of DCAE and GDCAE contained FA 16:0 (palmitic acid), 18:1 (putatively oleic acid), and 18:2 (putatively linoleic acid), the ratios of DCAE and GDCAE clearly correlated with the abundance of *Alistipes*. Since *Alistipes* and its phylum *Bacteroidota* are known to have bile acid hydrolytic (BSH) enzymatic activity, the ratio of conjugated to deconjugated bile acid esters may be attributable to *Bacteroidota*, including *Alistipes*^21^.

Likewise, we characterized additional three conjugated bile acid candidates, which include SE 24:1;O3;G/FA (glycine conjugated lithocholic acid ester, GLCAE, as a representative structure), SE 24:1;O4;T/FA (taurine conjugated deoxycholic acid ester, TDCAE, as a representative structure), and SE 24:1;O3;T/FA (taurine conjugated lithocholic acid ester, TLCAE, as a representative structure). These bile acid esters were validated by checking the diagnostic ions to recognize the aglycone and free fatty acid features (**Supplementary Table 8**). Furthermore, mannosylinositol phosphorylceramide (MIPC) was also characterized in the human cohort data because of the existence of phosphatidylinositol ceramide (PI-Cer) classification by the MS2lipid program, in addition to a unique neutral loss of hexose^22^. It is known that not only bacteria but also fungi exist in the human gut^23^. The detection of MIPC, a lipid produced by fungi, in a human cohort study was reasonable. Finally, we discovered phosphatidyl threonine (PT) in a mouse aging study, where the spectrum of PT was PS with a probability of 99.8%^24^. PT molecules were characterized in the kidneys and lungs of mice, but no statistical significance was observed with aging.

## Discussion

The MS2Lipid program has been developed to provide probability scores for lipid subclasses in MS/MS queries using a machine learning approach. It has been trained on over 10,000 lipid MS/MS spectral data and has shown accurate lipid subclass estimation with 95.5% accuracy. Compared to the CANOPUS metabolite ontology prediction program, MS2Lipid outperformed in terms of performance. Unlike MS-DIAL, which provides the most probable lipid candidate for a single query, MS2Lipid presents estimated lipid subclasses along with their probabilities. This feature is particularly useful for understanding mixed spectra of multiple co-eluting metabolites Additionally, we demonstrated the potential application of MS2Lipid for structural estimation of unknown spectra when used in combination with molecular spectrum networking.

The MS-DIAL program introduced in 2020 offers accurate and comprehensive lipid annotation using rule-based fragment ion annotations with predicted retention time information. However, the annotation results are often revised through manual curation by experienced technicians. The reasons why data curators modify the MS-DIAL results vary. For example, while fatty acids such as 18:4 or 18:5 typically do not exist in mammalian cells, complex lipids containing them are often suggested in the MS-DIAL software program. Such annotations of minor fatty acids may be true; however, conservative analysts often exclude these annotations. Moreover, even if a spectrum can be clearly interpreted, annotations are omitted if the chromatographic peak shape is poor or if the peak has a low signal-to-noise ratio, which makes quantitative discussion difficult. Similarly, when a product ion spectrum can be interpreted as a mixture of PC and PE, as demonstrated in this study, the peak feature is often excluded from the lipidome table in statistical analyses because the quantitative discussion becomes challenging. From 2020 to 2023, our research group has been labeling each lipid annotation result output by MS-DIAL with “confidence (correct),” “mixed spectrum,” “uncertainty,” and “misannotation (false hits)” labels, by using MS-DIAL’s graphical user interface (GUI). The training set used in this study only utilized those labeled with “confidence (correct),” reflecting high certainty, and moreover, it reflects the labeling results of just a single experienced analyst.

Machine learning from training sets derived from a single curator has both advantages and disadvantages. The advantage is the ability to maintain data quality. Additionally, it provides clear decision-making for spectral queries, although judgments may vary among individuals. In untargeted lipidomics, which is oriented towards discovery science, some researchers opt to curate spectra in a way that tolerates a certain level of false positives, aiming to reduce false negatives. Stearidonic acid, an omega-3 fatty acid derived from plant oils, is a representative candidate for 18:4 mentioned above, and lipids containing 18:4 can be detected in the human body^25^. The possible intake of such exposomes introduces ambiguity, leading to variations in data curation among analysts. In fact, our results in the MS2Lipid evaluation showed that differences among data curators had a greater impact on accuracy estimation than machine differences. Thus, decisions that reflect the will of an analyst can be interpreted as both strengths and weaknesses. This disadvantage may be due to the slow pace of spectral data accumulation. However, from the perspective of maintaining data quality, this seems to be a tradeoff. We plan to continue developing our approach, including quality control methods, as we accumulate more spectra for machine learning in the future.

Furthermore, the study investigated a newly characterized lipid subclass, GDCAE, in relation to microbiome data. GDCAE molecules were found to be significantly increased in obese individuals, which is associated with the decreased activity of BSH, a factor linked to obese-related diseases such as type-2 diabetes^26^. The abundances of DCAE and GDCAE were highly correlated with the abundances of the *Alistipes* genus, which has shown improvements in lipid accumulation and insulin resistance through the reduction of gut monosaccharide levels^12^. Understanding the impact of glycine-conjugated bile acid esters on human intestinal immune mechanisms and their relationship with bacteria, including *Alistipes*, may provide valuable insights for probiotics and prebiotics. MS2Lipid is not only a tool for lipid annotation but also a platform for gaining new insights into lipid biology. Future improvements to MS2Lipid aim to predict not only lipid subclasses but also more detailed structures, including fatty acid acyl chains, to contribute to the advancement of untargeted lipidomics.

## Supporting Information

## Supplementary Figures

**Supplementary Figure 1.**
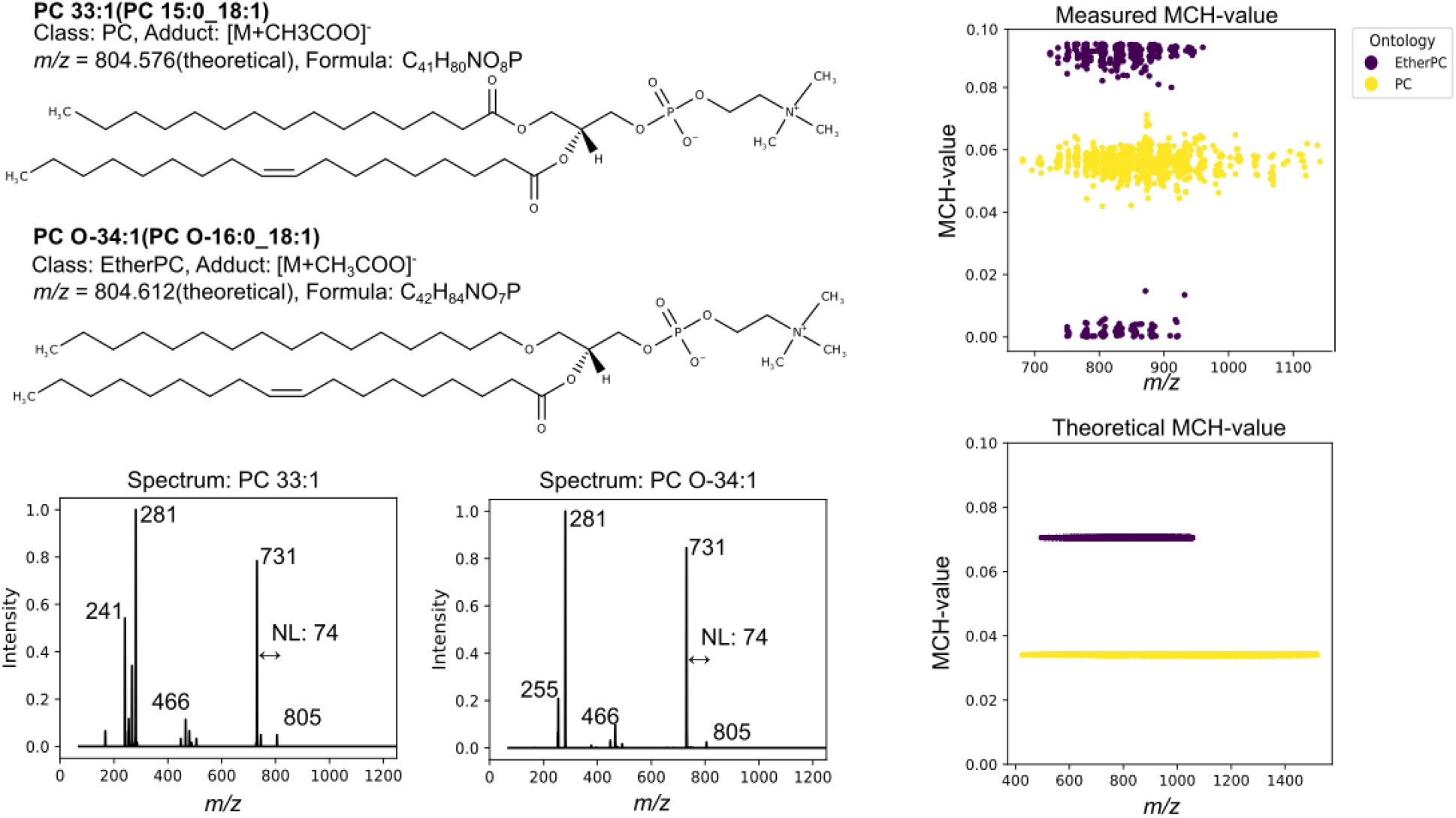
An example of MS/MS spectra and their MCH-values. Classification of MCH values based on PC and EtherPC structural formulas, MS/MS spectra, and oxygen atom counts in negative ion mode. The chemical structures and MS/MS spectra of PC 15:0_18:1 and PC O-16:0_18:1 were described. The MCH values of PC and ether PC included in our training set were calculated for the experimental (top-right panel)- and theoretical precursor (bottom-right panel) *m/z* values, respectively.

## Data availability

The training, validation, and test data for constructing the MS2lipid program are presented in Supplementary Data 2. The experimental MS/MS spectra for validation of MS2lipid were downloaded from the RIKEN Lipidomics website (http://prime.psc.riken.jp/menta.cgi/lipidomics/index). Human cohort lipidomics data were downloaded from the RIKEN DROPMet website (http://prime.psc.riken.jp/menta.cgi/prime/drop_index) under the index number DM0037. The source code and tutorial for MS2lipid are available at https://github.com/systemsomicslab/ms2lipid.

## Author contributions statement

H.T. and M.A. designed this study. N.S., T.O., Y.M., and K.N. developed MS2Lipid, and N.S. evaluated the program. A.H. curated the lipidomics data used for machine learning. N.S. analyzed the public data reanalysis. N. S. and H. T. wrote the manuscript. All authors have thoroughly discussed this project and helped improve the manuscript.

## Acknowledgements

This study represents a portion of the dissertation submitted by Nami Sakamoto to the Tokyo University of Agriculture and Technology in partial fulfillment of her Ph.D. requirements. This work was supported by the JST ERATO “Arita Lipidome Atlas Project” (JPMJER2101 to M.A. and H.T.), JSPS KAKENHI (20H00495 to M.A. and 24K02011 to H.T.), the National Cancer Center Research and Development Fund (2023-A-08, H.T.), AMED Moonshot Research and Development Program (JP22zf0127007 to M.A.), AMED NEDDTrim (22ae0121036h0002 to M.A.), AMED Japan Program for Infectious Diseases Research and Infrastructure (21wm0325036h0001, H.T. and M.A.), AMED Brain/MINDS (JP15dm0207001 to H.T.), JST National Bioscience Database Center (JPMJND2305 to H.T.), and Takeda Science Foundation (to M.A.). The funders had no role in the study design, data collection and analysis, decision to publish, or manuscript preparation.

