## Supplementary Note 1 for "MS2Lipid: a lipid subclass prediction program using machine learning and curated tandem mass spectral data"

Consider any two lipids, L_1_ and L_2_, that belong to the same lipid class differing only in carbon chain length and number of double bonds. The exact masses of L_1_ and L_2_ are denoted as M_L1_ and M_L2_ respectively. Here, the following relationship holds:

$$M_{L_{1}}=M_{L_{2}}+n_{1}CH_{2}+ n_{2}H_{2} \cdots(1)$$

CH_2_ and H_2_ are the exact masses of CH_2_ and H_2_, respectively, and n_1_ and n_2_ are arbitrary integers. Here, define the MCH-value for a lipid L as follows:

$$MCH_{L}=\left( \left( M_{L} mod CH_{2} \right) mod H_{2} \right) mod \left( 7H_{2}- CH_{2} \right) \cdots(2)$$

The MCH-values for L1 and L2 are:

$$MCH_{L_{1}}=\left( \left( M_{L_{1}} mod CH_{2} \right) mod H_{2} \right)mod \left( 7H_{2}- CH_{2} \right) \cdots(3)$$

$$MCH_{L_{2}}=\left( \left( M_{L_{2}} mod CH_{2} \right) mod H_{2} \right)mod \left( 7H_{2}- CH_{2} \right)\cdots(4)$$

From equation (1):

$$MCH_{L_{1}}=\left( \left( \left( M_{L_{2}}+n_{1}CH_{2}+n_{2}H_{2} \right) mod CH_{2} \right)mod H_{2} \right) mod \left( 7H_{2}- CH_{2} \right)$$

$$=\left( \left( \left( M_{L_{2}} mod CH_{2}+n_{2}H_{2} mod CH_{2} \right) mod CH_{2} \right) mod H_{2} \right) mod \left( 7H_{2}- CH_{2} \right)$$

$$=\left( \left( M_{L_{2}} mod CH_{2}+n_{2}H_{2} mod CH_{2}-r_{CH_{2}}CH_{2} \right) mod H_{2} \right) mod \left( 7H_{2}- CH_{2} \right)$$

$$=\left( \left( \left( M_{L_{2}} mod CH_{2} \right) mod H_{2}+\left( n_{2}H_{2} mod CH_{2} -r_{CH_{2}}CH_{2} \right) mod H_{2}- r_{H_{2}}H_{2} \right) mod H_{2} \right) mod \left( 7H_{2}- CH_{2} \right)$$

$$= \left( \left( M_{L_{2}} mod CH_{2} \right) mod H_{2}+{(n}_{2}H_{2} mod CH_{2} -r_{CH_{2}}CH_{2}) mod H_{2}- r_{H_{2}}H_{2} \right) mod \left( 7H_{2}- CH_{2} \right)$$

$$= \left( \left( \left( M_{L_{2}} mod CH_{2} \right) mod H_{2} \right) mod \left( 7H_{2}- CH_{2} \right)+\left( \left( n_{2}H_{2} mod CH_{2} -r_{CH_{2}}CH_{2} \right) mod H_{2}- r_{H_{2}}H_{2} \right) mod \left( 7H_{2}- CH_{2} \right) \right) mod \left( 7H_{2}- CH_{2} \right)$$

$$=\left( MCH_{L_{2}}+\left( \left( n_{2}H_{2} mod CH_{2} -r_{CH_{2}}CH_{2} \right) mod H_{2}- r_{H_{2}}H_{2} \right) \right) mod \left( 7H_{2}- CH_{2} \right) \cdots(5)$$

Where r_CH2_ and r_H2_ are defined as:

$$r_{CH_{2}}=\left\{ \begin{aligned} 1 (if M_{L_{2}} mod CH_{2}+n_{2}H_{2} mod CH_{2}>CH_{2}) \\ 0 (otherwise) \end{aligned} \right.$$

$$r_{H_{2}}= \left\{ \begin{aligned} 1 (if \left( M_{L_{2}} mod CH_{2} \right) mod H_{2}+\left( n_{2}H_{2} mod CH_{2} -r_{CH_{2}}CH_{2} \right) mod H_{2}>H_{2}) \\ 0 (otherwise) \end{aligned} \right.$$

Using integers $m, k$, let $n_{2}=7m+ k$, where $0\leq k<7$.

Since $kH_{2}<CH_{2}$, the second term of the first item in equation (5) is:

$$\left( \left( n_{2}H_{2} mod CH_{2} -r_{CH_{2}}CH_{2} \right) mod H_{2}- r_{H_{2}}H_{2} \right) mod \left( 7H_{2}- CH_{2} \right)$$

$$= \left( \left( n_{2}H_{2} mod CH_{2} mod H_{2} -r_{CH_{2}}CH_{2} \right) mod H_{2} - r_{H_{2}}H_{2} \right) mod \left( 7H_{2}- CH_{2} \right)$$

$$= \left( \left( \left( 7m+k \right)H_{2} mod CH_{2} mod H_{2} -r_{CH_{2}}CH_{2} \right) mod H_{2} - r_{H_{2}}H_{2} \right) mod \left( 7H_{2}- CH_{2} \right) \cdots(6)$$

Since $kH_{2}<CH_{2}$, $kH_{2} mod CH_{2} mod H_{2}=0$

Moreover, since $6H_{2}<CH_{2}<7H_{2}$, $7H_{2} mod CH_{2}<H_{2}$, thus:

$$\left( \left( 7m+k \right)H_{2} mod CH_{2} \right) mod H_{2}=7mH_{2} mod CH_{2}=\left( 7H_{2}-CH_{2} \right)m mod CH_{2} \cdots(7)$$

From equation (7), equation (6) is:

$$(6)= \left( \left( \left( 7H_{2}-CH_{2} \right)m mod CH_{2}-r_{CH_{2}}CH_{2} \right) mod H_{2} - r_{H_{2}}H_{2} \right) mod \left( 7H_{2}- CH_{2} \right)$$

$$= \left( \left( \left( 7H_{2}-CH_{2} \right)m mod CH_{2}+r_{CH_{2}}\left( -CH_{2} mod H_{2} \right) mod H_{2} \right) mod H_{2} - r_{H_{2}}H_{2} \right) mod \left( 7H_{2}- CH_{2} \right)$$

$$= \left( \left( \left( 7H_{2}-CH_{2} \right)m mod CH_{2}+r_{CH_{2}}\left( 7H_{2}-CH_{2} \right) \right) mod H_{2} - r_{H_{2}}H_{2} \right) mod \left( 7H_{2}- CH_{2} \right)$$

$$= \left( \left( \left( 7H_{2}-CH_{2} \right) \left( m mod CH_{2} \right)+r_{CH_{2}}\left( 7H_{2}-CH_{2} \right) \right) mod H_{2} - r_{H_{2}}H_{2} \right) mod \left( 7H_{2}- CH_{2} \right)$$

$$= \left( \left( \left( r_{CH_{2}}+m mod CH_{2} \right)\left( 7H_{2}-CH_{2} \right) \right) mod H_{2} - r_{H_{2}}H_{2} \right) mod \left( 7H_{2}- CH_{2} \right)$$

$$= \left( \left( r_{CH_{2}}+m mod CH_{2} \right)\left( 7H_{2}-CH_{2} \right) - r_{H_{2}}H_{2} \right) mod \left( 7H_{2}- CH_{2} \right)$$

$$= \left( -r_{H_{2}}H_{2} \right) mod \left( 7H_{2}- CH_{2} \right)$$

Thus, when $r_{H_{2}}=0$, $MCH_{L_{1}}=MCH_{L_{2}}.$

Therefore, for any two lipids L_1_ and L_2_ belonging to this lipid class, MCH_L1_ = MCH_L2_ holds.
